# Attempts to implement CRISPR/Cas9 for genome editing in the oomycete *Phytophthora infestans*

**DOI:** 10.1101/274829

**Authors:** Johan van den Hoogen, Francine Govers

## Abstract

Few techniques have revolutionized the molecular biology field as much as genome editing using CRISPR/Cas9. Recently, a CRISPR/Cas9 system has been developed for the oomycete *Phytophthora sojae*, and since then it has been employed in two other *Phytophthora* spp. Here, we report our progress on efforts to establish the system in the potato late blight pathogen *Phytophthora infestans*. Using the original constructs as developed for *P. sojae*, we did not obtain any transformants displaying a mutagenized target gene. We made several modifications to the CRISPR/Cas9 system to pinpoint the reason for failure and also explored the delivery of pre-assembled ribonucleoprotein complexes. With this report we summarize an extensive experimental effort pursuing the application of a CRISPR/Cas9 system for targeted mutagenesis in *P. infestans* and we conclude with suggestions for future directions.

## Introduction

The oomycete *Phytophthora infestans*, the causal agent of potato and tomato late blight, is an economically important plant pathogen that is difficult to control (Kamoun et al. 2015). Current efforts to identify and functionally analyze genes important for virulence, are hampered by a limited molecular toolbox. Gene knock-outs (KOs) or gene deletions are not possible via homologous recombination, as transgenes are integrated randomly via non-homologous end-joining (NHEJ) (Judelson 1997; Fang & Tyler 2016). Moreover, *P. infestans* is diploid and heterothallic. Generating sexual progeny is quite challenging and as a result studies based on genetic analyses are scarce (Govers & Gijzen 2006; Fry 2008). Consequently, to date, functional gene studies in oomycetes primarily rely on gene silencing and overexpression. DNA transformation of the target gene in *P. infestans* can lead to homology dependent gene silencing or overexpression (van West et al. 1999). By fusing a fluorescent tag to the target gene, the overexpression transformants can also be exploited for investigating the subcellular localization of the encoded protein. However, due to the random integration of transgenes and varying levels of silencing and overexpression efficiency, phenotypes often vary between transformed lines, experiments, and labs (Fang & Tyler 2016).

Recently, a CRISPR/Cas9 genome editing system has been developed for *Phytophthora sojae* (Fang & Tyler 2016). This provides an important addition to the molecular toolbox for oomycetes, and makes it up to par with other research areas. For genome editing based on CRISPR/Cas systems, a nuclease, e.g. Cas9, is targeted to the desired DNA sequence by a so-called synthetic guide RNA (gRNA). The 20-nucleotide protospacer directs Cas9 to a specific target DNA site, which must be immediately 5’ of a protospacer adjacent motif (PAM) sequence (Ran et al. 2013). When Cas9 is directed to the target site a double-stranded break (DSB) is induced, which can be repaired either by the NHEJ repair mechanism, or via homology-directed repair (HDR) in case a repair template is available (Ran et al. 2013). Because of its low fidelity, NHEJ often leads to insertions or deletions (indels) in the repaired target gene, which can lead to frameshift mutations. HDR, on the other hand, can be exploited to introduce specific modifications such as base substitutions, insertions, or gene deletions.

So far, the *P. sojae* CRISPR/Cas9 system has been successfully used to mutagenize several genes in *P. sojae* (Fang & Tyler 2016; Ma et al. 2017), *Phytophthora capsici* (B.M. Tyler, personal communication), and *Phytophthora palmivora* (Gumtow et al. 2017). However, as of to date, no one has reported successful implementation of the system in *P. infestans*. Here, we present a case study for the effectuation of CRISPR/Cas9 for targeted genome editing in *P. infestans*. We chose to target three genes in *P. infestans* for a proof-of-principle study of CRISPR/Cas9-based genome editing in this organism.

The first target, *Avr1*, encodes an RXLR effector protein recognized by the cognate receptor protein R1 in potato. Recognition of AVR1 by R1 provides resistance to potato, while *P. infestans* strains lacking *Avr1* are virulent on potato cultivars harboring the resistance gene *R1*. Moreover, R1 fails to recognize several mutated forms of AVR1 (Du et al. 2018). Hence, *P. infestans Avr1* KO lines are expected to gain virulence on *R1* potato, providing a clear phenotype compared to non-edited lines. The second target, *PiTubA2*, is one of the five α-tubulin encoding genes in *P. infestans*. Our aim was to modify the endogenous *PiTub2A* gene in such a way that it encodes a fusion protein that has a GFP tag fused to the N-terminus of PiTUBA2. We therefore designed a HDR construct for gene knock-in at the endogenous *PiTubA2* locus. A PiTubA2-GFP knock-in line would be ideal for live cell imaging of the microtubule cytoskeleton in *P. infestans*, as it lacks possible non-desirable side effects associated with overexpression and/or ectopic integration of the transgene. The third target, *PiAP5*, encodes an aspartic protease (AP) with an C-terminal GPCR domain (Kay et al. 2011), a protein which is unique for oomycetes (van den Hoogen et al. 2018). To study to what extent the GPCR domain is important for functioning of the AP domain, we aimed at targeting Cas9 to the central region of the gene that separates the parts encoding the AP domain and the GPCR domain. An indel in this region should result in a truncated protein, which contains the N-terminal AP domain but lacks the C-terminal GPCR domain.

In the initial setup of our experiments, we used the CRISPR/Cas9 system as developed by Fang and Tyler (2016). In this system, the expression of Cas9 is driven by the strong constitutive *Bremia lactucae* Ham34 promoter and the expression of the gRNA by the *P. sojae* RPL41 promoter. The RPL41 promoter was chosen because the U6 small nuclear RNA promoters typically used in CRISPR/Cas9 systems, did not yield any detectable transcript (Fang et al. 2017). To ensure correct release of the gRNA from the transcript, the gRNA sequence was flanked with a hammerhead and a hepatitis delta virus (HDV) ribozyme at the 5’ and 3’ side, respectively (Fang et al. 2017).

This report summarizes an extensive experimental effort pursuing the application of a CRISPR/Cas9 system for targeted mutagenesis in *P. infestans* and concludes with suggestions for future directions.

## Results and discussion

### Experimental setup: design of gRNAs

The first, and perhaps most critical step of setting up a CRISPR/Cas9 experiment, is the design of the gRNAs. The genes of interest were screened for target sites using NGG as the PAM. All gRNAs were scored for on-target activity according to Doench *et al*. (Doench et al. 2014) and for off-targets interactions according to Hsu *et al*. (Hsu et al. 2013). The top 10 best scoring proto-spacers were manually curated using BLAST analysis on the *P. infestans* reference genome and predictions of the secondary RNA structure of the corresponding gRNAs. For *Avr1*, we selected the three best scoring gRNAs (**Figure 1a**), i.e. no predicted off-target interactions and no strong secondary RNA structure. For *PiTubA2* and *PiAP5* we selected the highest scoring gRNA (**Figure 1b, 1c**).

**Figure 1.**
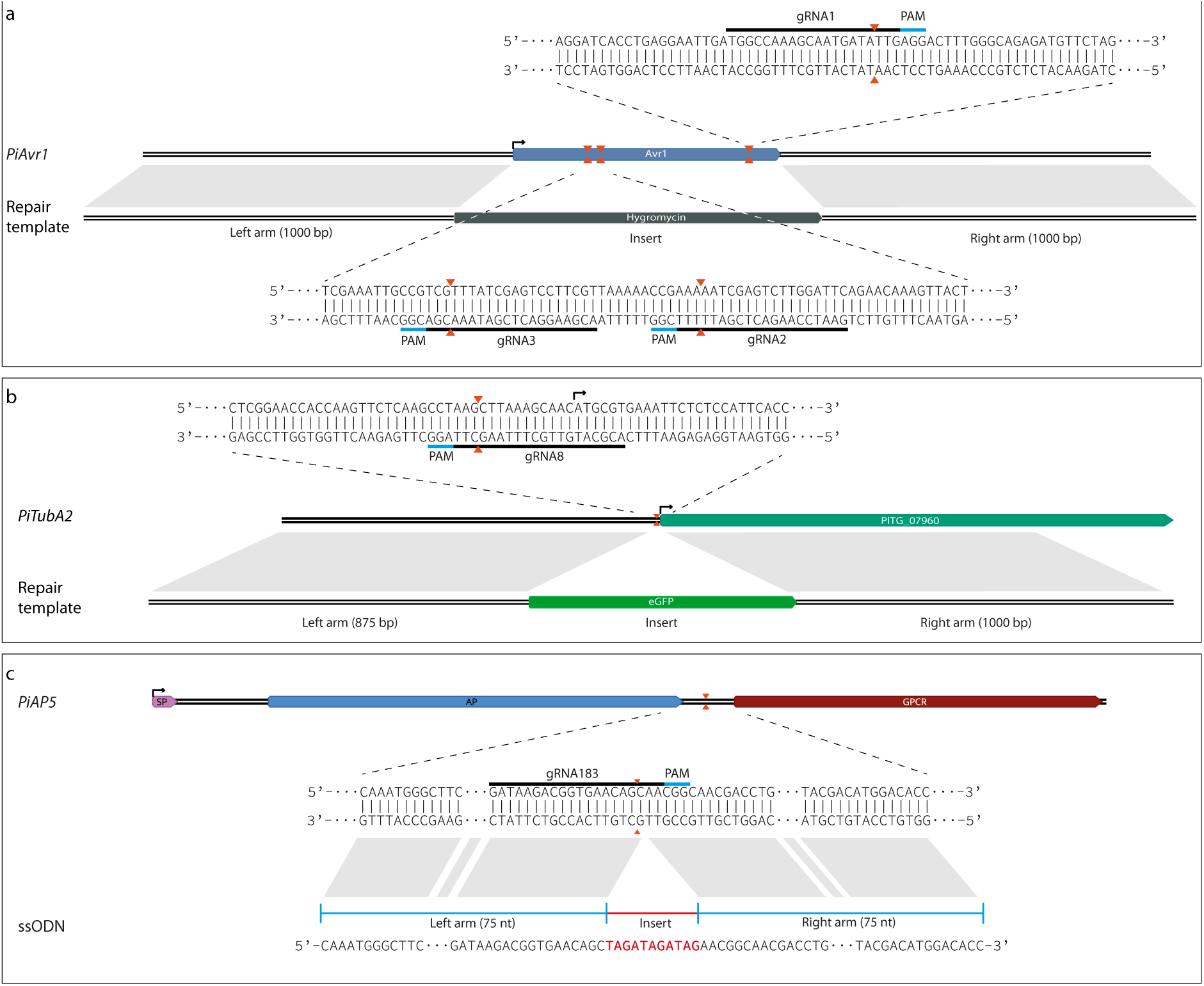
CRISPR loci and HDR constructs. **(a)** *PiAvr1*, **(b)** *PiTubA1*, and **(c)** *PiAP5*. Orange arrowheads indicate expected DSB sites upon Cas9 nuclease activity; black arrows indicate start codons (ATG); grey blocks mark homologous regions between the genes and the HDR constructs (referred to as repair template or ssODN); interpuncts (•) represent cropped sequences; PAM: Protospacer Adjacent Motif.

### HDR constructs

To employ the HDR repair pathway for targeted mutagenesis, we used a similar approach as previously used in *P. sojae*. This implies cotransformations of three plasmids, i.e., two plasmids with sequences encoding Cas9 and gRNA, respectively, and the third one containing the HDR repair template (Fang & Tyler 2016).

For *Avr1* we designed a construct to replace the coding region with *HygB*, a hygromycin-B resistance gene (**Figure 1a**). The inserts were flanked by a 1 kb right flanking arm and a 875 bp left flanking arm that are complementary to the target site for recombination. The reason for the shorter left flanking arm is a 380 bp region in the genome assembly, that is not accessible for sequencing. We opted to use the largest possible flanking region, i.e. 875 bp (**Figure 1a**). For *PiTubA2* we designed a construct to knock-in GFP just before the start codon of the open reading frame (**Figure 1b**). The repair template was designed as such that the PAM would be disrupted upon HDR repair.

For a number of organisms, higher efficiency of HDR repair has been reported when making use of single-stranded oligodeoxynucleotides (ssODN) as repair template, instead of plasmid DNA (Chen et al. 2011). We designed a ssODN repair template for *AP5*, comprising an 11 bp insert flanked by two 75 nt homology arms. The insert is designed in such a way that there is a stop codon (TAG) in each of the three open reading frames (**Figure 1c**). Further, the ssODN and gRNA were designed as such, that introduction of the 11 bp construct would disrupt the protospacer and consequently withhold Cas9 from further processing the target site. The expected protein product from the truncated gene contains the AP domain but lacks the GPCR domain.

### Testing the detection limit

We next examined the sensitivity of our screening methods. To do so, we simulated the situation that an unknown fraction of transformants contains a Cas9-induced mutation. This is expected to be the case in a sample from pooled transformants after transformation; screening these for mutations can give an idea about the efficiency of targeted mutagenesis. First, we constructed an *Avr1* amplicon (ΔAvr1) containing the 29 bp deletion that is expected after Cas9 cleavage at sites gRNA2 and gRNA3 (**Figure S1a**). Next, varying molar ratios of *Avr1* and ΔAvr1 amplicons were annealed and incubated with T7 endonuclease I (T7EI), which is an enzyme that specifically digests mispaired DNA. We found that the detection limit is just over a molar ratio of 95:5 (i.e. 5% mutated amplicons) (**Figure S1b**).

In parallel, PCR amplicons were sequenced. When sequencing an amplicon that is obtained from a PCR on a pool of transformants of which a certain fraction underwent CRISPR-induced mutagenesis, the resulting sequence chromatogram will contain a certain amount of ‘noise’, or background signals. These aberrant chromatogram peaks reflect the approximate frequency of amplicons with an indel, which can be estimated with an analysis tool such as TIDE (Brinkman et al. 2014). TIDE quantifies the editing efficacy by calculating the statistical probability of finding background signals in the chromatogram of a sample compared to finding them in a reference sequence chromatogram. A range of molar ratios of *Avr1*:ΔAvr1 was sequenced, and by using T30-2 *Avr1* as a reference, we analyzed the sequence chromatograms for mismatches. We found the lowest statistically significant detection at a molar ratio of 998:2 (i.e. 0.2% of the sequences having the deletion) (**Figure S1c**).

These detection limits are well below the observed frequencies of CRISPR-induced mutations in *P. sojae*, where up to 80% of the analyzed transformants have undergone NHEJ (Y.F. Fang, personal communication). Assuming that the CRISPR/Cas9 system is functional in *P. infestans*, our detection assays are sufficiently sensitive to reveal induced mutations. It should be noted however, that in the simulations described above only a single mutation (i.e. a 29 bp deletion) was present. In an actual sample of pooled transformed *P. infestans* protoplasts, a mix of differently sized indels is expected and the detection limit will be higher. Nonetheless, we assume that our detection assays are sufficiently sensitive to detect CRISPR-induced mutations in pooled *P. infestans* transformed protoplasts.

### *In vitro* activity assay

To examine the functionality of the designed gRNAs, we performed an *in vitro* cleavage assay of target DNA. This assay can give an indication of the *in vivo* efficacy of the gRNA for Cas9 activity. We used purified Cas9 protein and *in vitro* transcribed gRNA in equimolar amounts, in combination with a PCR product as target DNA. We observed Cas9 activity on *Avr1* using gRNA1 and gRNA3 (**Figure 2a**). In contrast, gRNA2 failed to direct Cas9 to the target site (**Figure 2a**). Examination of the sequence revealed a mistake in the design of the initial construct. When corrected, gRNA2 also showed activity (not shown). We did not observe Cas9 activity on *PiTubA2* using gRNA8, nor on *PiAP5* using gRNA183 (**Figure 2b**). In contrast to the initial gRNA2 on *Avr1*, we could not detect mistakes in the design of these two gRNAs. Consequently, judging from their inability to guide Cas9 *in vitro*, it is likely that also the *in vivo* efficacies of gRNA8 and gRNA183 are limited. Other gRNAs for these target genes may have an increased activity, but so far we have not tested alternative gRNAs for *PiTubA2* and *PiAP5*.

**Figure 2.**
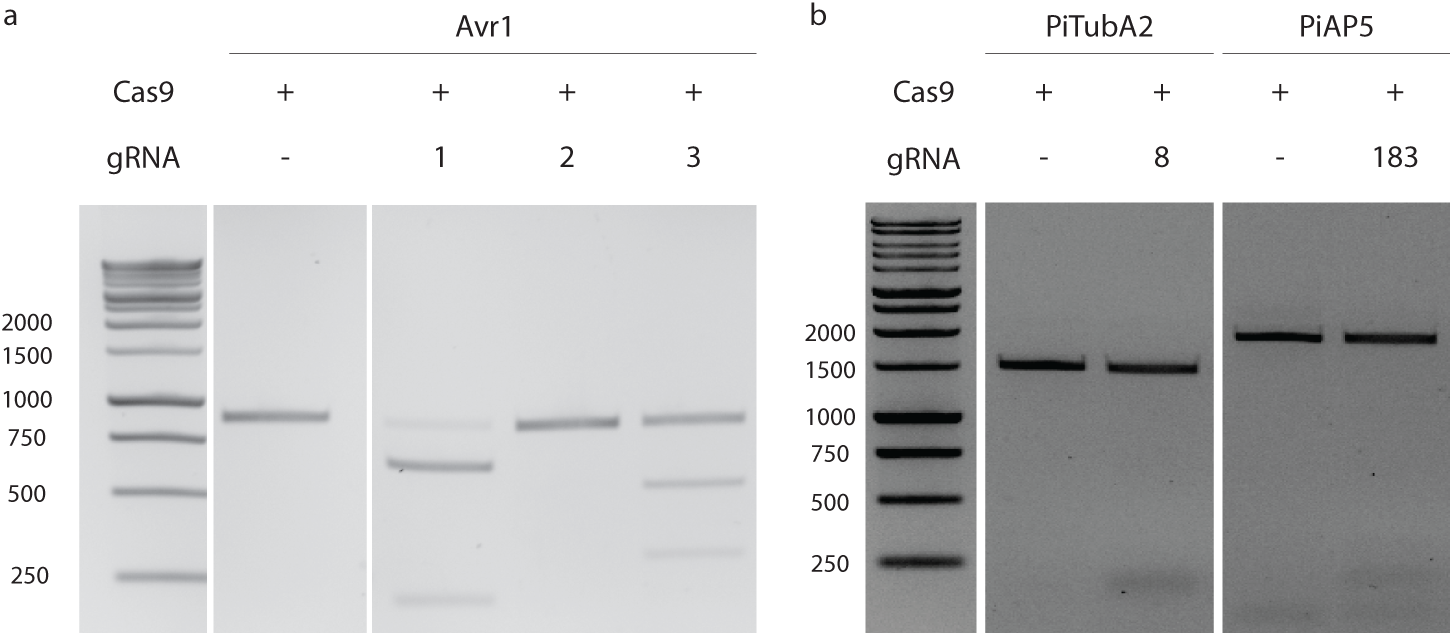
In vivo cleavage assay. **a)** gRNAs 1, 2, and 3 with Avr1 as target DNA. **b)** gRNA8 with PiTubA2 as target DNA (**right**) and gRNA183 with PiAP5 as target DNA (**left**).

### *In vivo* CRISPR*/*Cas9 in *P. infestans*

Next, we set out to test the *in vivo* editing efficacy of the system. To this end, we first cloned the respective gRNAs into the expression plasmid pYF2.3-gRNA (Fang & Tyler 2016). We performed several transformations of *P. infestans* with plasmids pYF2.2-Cas9 and pYF2.3-gRNA and obtained numerous transformants resistant to the selection antibiotic geneticin. In these transformants, the presence of indels as a result of the presumed Cas9 activity was monitored by T7EI digestion and sequencing analysis. However, contrary to expectations set by the positive results of the use of CRISPR/Cas9 in *P. sojae, P. capsici*, and *P. palmivora*, the analyses did not reveal any lines in which the target gene was mutagenized. In some cases, the sequence chromatogram showed ambiguities near the target site, raising the hope that the mutagenesis was successful. More detailed analysis however, revealed that these lines were identical to the recipient isolate and should be considered as false-positives. Further trials focused at targeting *Avr1* included transformations combining two or three gRNA-expressing vectors along with Cas9, but in all cases the results were negative.

In order to induce specific mutations in *P. infestans*, we performed experiments where plasmid DNA containing a HDR repair template for *Avr1* was co-transformed with the CRISPR/Cas9 constructs. Even though several hygromycin-B resistant colonies were obtained, and the presence of *HygB* could be detected by PCR on genomic DNA, we were not able to detect integration of the transgene at the endogenous genomic locus. Possibly, the hygromycin-B resistance gene was integrated ectopically, or resistance was conferred by transcription from non-integrated plasmid DNA. Similarly, we did not obtain transformed lines with GFP-encoding sequences inserted at the 5’ end of *PitubA2*, or *PiAP5*-mutants with the stop codon insertion.

For the target genes *PiTubA1* and *PiAP5* only a single gRNA was used. This and the fact the both gRNAs did not prove to be effective in the *in vitro* activity assay, are most likely the reasons for the absence of observable Cas9 activity. However, also in the case of *Avr1* as target gene, where all three gRNAs proved functional *in vitro*, Cas9 activity appeared to be absent.

An obvious explanation for failure is poor expression of Cas9. To assess the expression level of Cas9, reverse-transcription PCR was performed on RNA isolated from mycelium of transformed lines. In several lines expression of Cas9 could be detected (**Figure S2**). Consequently, it is unlikely that lack of expression of Cas9 is the prime reason for failure of the system. As yet, we have not monitored the presence of the gRNA transcripts. Altogether, these findings led us to conclude that the CRISPR/Cas9 system in *P. infestans* is not functioning, at least not up to the expected efficiency.

### Modifications to the CRISPR*/*Cas9 system

It is hard to pinpoint the main cause for failure of the system, as multiple factors may have a role. Hence, we set out to make modifications at different steps in the process. Below we describe several modifications and alternative approaches in an effort to effectuate the CRISPR/Cas9 system in *P. infestans*.

#### Alternative vectors

In the original *P. sojae* CRISPR/Cas9 system, there is no selectable resistance marker present on the plasmid encoding the gRNA. However, in practice, co-transformations of plasmids usually result in integration of both plasmids (R. Weide, personal communication). Consequently, geneticin-resistant colonies are expected to also have incorporated the gRNA plasmid. Nevertheless, we decided to test whether we would be able to observe CRISPR/Cas9 activity by using a gRNA-encoding plasmid containing *HygB* as a resistance marker. To do so, we introduced the *HygB* in pYF2.3-gRNA-Ribo, replacing *eGFP* from the vector and yielding pJH2.4-gRNA-Ribo-Hyg (**Figure S3**). In this vector backbone, the respective gRNAs for the three target genes were cloned. Along with the Cas9 encoding plasmid, the resulting plasmids were used for cotransformations of *P. infestans*. Even though we obtained numerous hygromycin-B-resistant tranformants, in none of the screened transformants we could detect mutagenized target genes.

In another attempt, we cloned the gRNA expression components in pYF2.2-Cas9, resulting in the ‘all-in-one’ vector pJH2.5-Cas9-gRNA (**Figure S3**). Numerous geneticin-resistant transformants were obtained and PCR analysis confirmed the presence of the Cas9 and gRNA genes. Unfortunately, no mutagenized target genes were observed. In addition, we constructed a vector containing both gRNA2 and gRNA3 (both targeting *Avr1*), but also here we did not obtain transformed lines with the expected deletion between the two target sites.

#### Nuclear localization sequence

Nuclear localization of Cas9 is essential. Oomycetes contain distinct nuclear localization sequence (NLS) signals, and commonly used mammalian NLS signals are not efficient in *P. sojae* (Fang & Tyler 2016; Fang et al. 2017). To overcome this, Fang & Tyler (2016) fused a synthetic NLS derived from a *P. sojae* bZIP transcription factor (TF) to Cas9. This NLS, further referred to as PsNLS, showed strong nuclear localization in *P. sojae* (Fang et al. 2017), and it is expected that it will perform likewise in *P. infestans*. To test the nuclear localization of Cas9, we obtained several transformed *P. infestans* lines carrying a PsNLS-Cas9-GFP fusion construct but unfortunately, for unknown reasons none of these lines showed fluorescence. Hence, we were unable to determine the localization of Cas9 in *P. infestans*. Anticipating that PsNLS is not able to target Cas9 to the nucleus in *P. infestans*, we set out to test a different NLS. Using a BLAST search, we identified the *P. infestans* homolog of the *P. sojae* bZIP TF, from which the NLS was obtained. This gene, PITG_11668, encodes a TF that was previously shown to have strong nuclear localization in *P. infestans* (Gamboa-Melendez et al. 2013). Next, we replaced the *P. sojae* NLS in vectors pYF2.3-PsNLS-Cas9 and pYF2.3-PsNLS-Cas9-GFP with the NLS region of PITG_11668 (PiNLS) to obtain the vectors pJH2.6-PiNLS-Cas9 and pJH2.6-PiNLS-Cas9-GFP, respectively (**Figure S3**). In addition, PiNLS was cloned N-terminally of GFP in the basic expression vector pGFP-N (Ah-Fong & Judelson 2011), to obtain pGFP-PiNLS-GFP (**Figure S3**). Next, these constructs were used for transformation of *P. infestans*. We performed cotransformations of pJH2.6-PiNLS-Cas9 along with gRNAs for *Avr1, PiTubA2*, or *PiAP5* carried by vectors pYF2.3-gRNA or pJH2.4-gRNA-Hyg. Unfortunately, for none of the combinations we did obtain transformed lines in which we could observe Cas9 activity, nor did we obtain transformed lines exhibiting a fluorescent signal.

#### gRNA expression

In the *P. sojae* CRISPR/Cas9 system, expression of the ribozyme-gRNA construct as well as that of *nptII*, the gene providing geneticin resistance, is driven by p*Ps*RPL41, the promoter of the *P. sojae* ribosomal gene *Ps*RPL41. Thus, geneticin-resistant transformants should in principle express the ribozyme-gRNA construct. However, in the numerous geneticin-resistant transformants that we have obtained, we did not observe CRISPR/Cas9 activity. Consequently, we questioned whether the promoter of *Ps*RPL41 is active in *P. infestans*. In an earlier study performed in *P. infestans*, the expression stability of the *P. infestans* ortholog of p*Ps*RPL41 was found to be inferior to other *P. infestans* and *P. capsici* ribosomal promoters (Poidevin et al. 2015). p*Ps*RPL41 was not evaluated. To assess whether the use of a different promoter driving the expression of the ribozyme-gRNA construct could improve CRISPR/Cas9 activity, we opted to use the *P. capsici* S9 promoter (p*Pc*S9). This promoter was found to provide high and stable expression in *P. infestans* (Poidevin et al. 2015). To replace p*Ps*RPL41 with p*Pc*S9, the promotor was PCR amplified from pTOR-S9 (Poidevin et al. 2015) and cloned into EcoRI/NheI sites of pYF2.3-gRNA-Ribo. However, an unexpected NheI site at approximately 150 bp upstream of the 3’ end of the promoter sequence, gave rise to a truncated insert, as the same restriction enzyme is used for gRNA insertion. To circumvent this problem p*Pc*S9 and the gRNA can be introduced into the vector backbone by Gibson assembly but due to time constraints we have not been able to obtain a suitable construct with p*Pc*S9 for transformation of *P. infestans*.

#### Delivery of ribonucleoprotein (RNP) complexes

In several organisms it has been shown that the delivery of Cas9 protein-gRNA ribonucleoproteins (RNPs) complexes is an efficient method for inducing targeted genome editing events, including a number of plant species (Woo et al. 2015), the nematode *Caenorhabditis elegans* (Cho et al. 2013), filamentous fungi and yeast (Pohl et al. 2016; Grahl et al. 2017), protozoa (Soares Medeiros et al. 2017), and also in human cell lines (Ramakrishna et al. 2014). Moreover, it was found to result in genome editing with substantially higher specificity compared to DNA transformation (Zuris et al. 2015). From a technical perspective, the use of RNP complexes may have several advantages over a ‘regular’ CRISPR system where both components are delivered as plasmid DNA and integrated into the genome. Firstly, the genome-editing efficiency of Cas9 using RNP complexes does not rely on transcription rate in the host, as both components are preassembled *in vitro* prior to delivery into cells. In *Phytophthora* spp., integration of foreign DNA occurs randomly, and it is plausible that integration in a ‘silent’ part of the genome results in reduced transcription rates. The integration can also lead to disruption of genes, when the transgene is inserted in an open reading frame. Secondly, and more importantly, transgene expression of Cas9 can have adverse effects and can even lead to cell death due to toxicity (Jiang et al. 2014; Kim et al. 2014; Peng et al. 2014). Generating CRISPR-induced mutations using RNP complexes might (partially) alleviate this issue, as after some time the complexes are degraded, reducing chances for side effects.

To our knowledge, the use of RNP complexes for targeted genome editing has not been explored in oomycetes. To assess whether RNP complexes can be used for targeted genome editing in *P. infestans*, we used a modified protocol for PEG-mediated protoplast transformation, substituting plasmid DNA with pre-assembled RNP complexes. In parallel, one sample was co-transfected with the *Avr1* HDR construct (**Figure 1a**), which has *HygB* as a selectable marker. After transfection, regenerated protoplasts were analyzed by a T7EI assay and sequencing. These analyses showed all samples to be identical to control transformants, with no significantly overrepresented aberrant background signals in the sequence chromatograms. Moreover, we could not confirm the integration of *HygB* at the target locus, not even in transformants that were hygromycin-B resistant.

The experimental setup of direct delivery of RNP complexes has some limitations, most importantly the lack of selection. Whereas in a regular plasmid-based transformation experiment the introduced plasmid contains a resistance gene to a selection antibiotic, protein transfection does not yield antibiotic resistant colonies and hence also non-transfected protoplasts will regenerate. Consequently, in our experiments transfected protoplasts might have been overshadowed by non-transfected protoplasts. Moreover, despite careful preparation of the starting material, some residual sporangia may have been present in the protoplast suspension and these also readily overgrow regenerating protoplasts.

Another potential limitation is the nuclear localization of the RNP complexes. The Cas9 protein used in this experiment contains two Simian virus 40 (SV40) T antigen NLS tags, one at the N-terminus and the other at the C-terminus. The SV40 NLS is a well-studied monopartite NLS tag consisting of several basic amino acids (Lange et al. 2007), and has been shown to localize fusion gene products to the nucleus of *P. sojae*, albeit with a reduced efficiency compared to other NLS tags (Fang et al. 2017). Hence, it is expected that the RNP complexes used in this study are localized to the nucleus. Still, the use of a Cas9 protein equipped with a *Phytophthora* specific NLS tag might increase the efficiency (Fang et al. 2017). Since such Cas9 proteins are not commercially available they would have to be produced in-house.

## Conclusions and future outlook

Contrary to expectations set by the successful application of the CRISPR/Cas9 system for targeted genome editing in *P. sojae, P. capsici* (B.M. Tyler, personal communication), and *P. palmivora* (Gumtow et al. 2017), we have as yet not been able to implement the system in *P. infestans*. The same holds for colleagues elsewhere who are also experiencing difficulties in effectuating the CRISPR/Cas9 system for use in *P. infestans* (personal communication). It is, however, hard to pinpoint the cause for failure of the system. Likely, it is an additive effect of several suboptimal conditions, such as Cas9 or gRNA expression levels, Cas9 localization, or the incubation temperature. Fang and Tyler (2016) who established the system in *P. sojae* also faced numerous challenges and made substantial modifications in the initial CRISPR/Cas9 procedure to get it to work in *P. sojae*. For *P. palmivora* Gumtow et al. (2017) choose *Agrobacterium-*mediated transformation (AMT) to implement the CRISPR/Cas9 system. Whereas protoplast transformation typically results in multiple integration events of the transgene(s), AMT usually gives rise to only one or two integrations of the transgene in the genome (Vijn & Govers 2003). A higher transgene copy number easily results in altered expression levels due to homology-dependent gene silencing or overexpression of the transgene. However, *P. infestans* transformants resulting from AMT rarely show silencing or overexpression of the target gene (P.J.I. van de Vondervoort, unpublished data), a phenomenon that might be due to the low integration rate. Overall, generating transformants via AMT is less laborious and requires less starting materials than PEG mediated protoplast transformations. We are currently exploring the use of AMT for integration of the CRISPR/Cas9 components in *P. infestans*.

A clear difference between *P. infestans* and the three *Phytophthora* spp. in which CRISPR-induced mutations are observed, is their growth temperature. Whereas *P. infestans* is typically incubated at 18°C, the other three species are grown at 25°C. The Cas9 isoform used in the *P. sojae* CRISPR/Cas9 system is a human codon optimized gene from *Streptococcus pyogenes* (SpCas9), an organism which grows at 37°C. Hence, it is anticipated that the activity of SpCas9 is reduced at decreased temperatures. Indeed, *Arabidopsis thaliana* plants exposed to heat stress (37°C for 30 h) showed much higher rates of targeted mutagenesis by CRISPR/Cas9 compared to plants grown continuously at the standard temperature of 22°C (Le Blanc et al. 2017). On the other hand CRISPR/Cas9 has been employed in salmon eggs at temperatures as low as 6°C (Edvardsen et al. 2014), making it unlikely that the incubation temperature is the sole reason for absence of Cas9 activity in *P. infestans*. Still, incubating potential *P. infestans* transformants at elevated temperatures or exposing them to heat stress, might improve the efficiency of the system and is worth to try. This would require some pilot experiments for determining the right temperature and time of exposure to the heat shock. When *P. infestans* is cultivated at 28°C it dies within 24 hours, but possibly a shorter exposure at elevated temperatures is sufficient to obtain higher frequencies of CRISPR-induced mutations.

Another possible modification to the system which might be considered, is the use of a different nuclease. *S. pyogenes* Cas9 (SpCas9) is a relatively large protein which might hamper nuclear import. For smaller isoforms, such as Cas9 from *Staphylococcus aureus* (SaCas9) with approximately three quarters of the size of SpCas9 (Ran et al. 2015), it might be easier to find its way to the nucleus. In the protozoa *Trypanosoma cruzi*, delivery of RNP complexes of SaCas9 and gRNA resulted in gene edits, while SpCas9 did not (Soares Medeiros et al. 2017). Likewise, RNP complexes with SaCas9 could be more effective in *P. infestans* compared to those with SpCas9. Also for plasmid-based delivery of the CRISPR/Cas9 components, the smaller gene size of *SaCas9* might improve integration. However, SaCas9 recognizes a different PAM than SpCas9 (NNGRRT vs. NGG). Hence, different gRNAs have to be designed when utilizing this nuclease, and there is likely a lower frequency of the PAM in the sequence of the target gene. Another nuclease to consider is Cpf1 (also known as Cas12a). Whereas SpCas9 creates DSB breaks with ‘blunt’ ends, Cpf1 introduces a staggered DSB break with a 4 or 5 nt 5’ overhang (Zetsche et al. 2015). These cohesive (‘sticky’) ends can increase the frequency of HDR repair. Moreover, on certain positions, Cpf1 is sensitive to single-base mismatches in the protospacer, reducing the frequency of off-target cleavage (Kleinstiver et al. 2016). Also, its T-rich PAM (TTTN) may limit the number of target sites in *P. infestans*, which has a GC content of 51% (Haas et al. 2009).

We trust that, with dedicated effort, developing a CRISPR/Cas system for *P. infestans* is attainable. Future work should focus on systematic analysis of factors limiting the efficiency of the system. When these limitations are identified and overcome, targeted mutagenesis in *P. infestans* might be within reach.

## Acknowledgements

We are grateful to Yufeng ‘Francis’ Fang and Brett Tyler for generously sharing constructs and experiences. Thomas van Ravensteyn for helpful discussions. Juliette Silven, Paul Schaap, and Natalie Verbeek-de Kruif are acknowledged for their technical assistance. This work was supported by Division for Earth and Live Sciences (ALW) with financial aid from the Netherlands Organization for Scientific Research (NWO) in the framework of the ALW-JSTP programme (project # 833.13.002; J.H.; F.G.)

## Materials and methods

### Strains, culture conditions and transformations

*P. infestans* strain T30-2 (van der Lee et al. 2001) and all transgenic lines were routinely grown at 18 °C in the dark on rye agar medium supplemented with 2% sucrose (RSA) (Caten & Jinks 1968). RSA was supplemented with 20 µg/ml vancomycin, 100 µg/ml ampicillin and 50 µg/ml amphotericin B, and in addition, for transformed lines with 2.5 µg/ml G418. Transient and stable transformants of *P. infestans* were generated using PEG/CaCl2-mediated protoplast transformation and zoospore electroporation. Protoplast transformation was performed according to methods described previously (Ah-Fong et al. 2008), omitting the step of complexing circular plasmid DNA with Lipofectin in protoplast transformation. Zoospore electroporation was performed following available protocols.

### gRNA design

Sequences for target genes were downloaded from FungiDB (Stajich et al. 2012). The full-length gene products were PCR amplified and subsequently validated by sequencing (Eurofins, Ebersberg, Germany). Next, the sequence was screened for CRISPR sites using NGG as PAM on Geneious R9.1.4 (Kearse et al. 2012). The resulting sites were scored for on-target activity according to Doench et al. (Doench et al. 2014), and for off-target interactions according to Hsu et al. (2013). The top 10 best scoring CRISPR sites were manually curated using BLAST analysis on the *P. infestans* reference genome and predictions of secondary RNA structure of the corresponding gRNAs using RNAstructure (Reuter & Mathews 2010).

gRNAs for *Avr1* and *PiTubA1* were ordered as sense and antisense PAGE-purified oligos (Integrated DNA Technologies) and subsequently annealed and ligated into pYF2.3-gRNA-Ribo-EV, as described by Fang and Tyler (2016), using BsaI and NheI-HF (New England Biolabs). For *PiAP5*, 283 bp constructs transcribing the ribozyme-gRNA insert flanked by two 30 bp overlaps, were designed and ordered as gBlocks (Integrated DNA Technologies). Next, the fragments were inserted into pYF2.3-gRNA-Ribo-EV by Gibson assembly using the NEBuilder HiFi kit (New England Biolabs Inc).

### Constructs

**Figure S3** shows the constructs used in this study. The hygromycin-B resistance gene was amplified from pGFP-H (Ah-Fong & Judelson 2011) using primers Hyg_AflII_F and Hyg_ApaI_R, and the resulting amplicon was cloned into pYF2.3-gRNA-Ribo using AflI and ApaI (New England Biolabs), yielding pJH2.4-Hyg-gRNA. To constitute the ‘all-in-one’ vector pJH2.5-Cas9-gRNA, the gRNA expressing construct from pYF2.3-gRNA (containing the RPL41 promoter, gRNA insert, and Hsp70 terminator) was inserted into pYF2.2-Cas9 using EcoRI (Promega). A dual-gRNA vector encoding Avr1-targeting gRNA2 and gRNA3 was constructed by inserting the ribozyme-gRNA3 construct into pYF2.3-gRNA2 by Gibson assembly. The amplicon containing the insert was made using primers gRNA3_Gibson_F and gRNA3_Gibson_R. The NLS region from PITG_11668 was inserted at the N-terminus of GFP, the PCR amplicon was cloned into the AgeI and NheI sites of pGFP-N (Ah-Fong & Judelson 2011), yielding pGFP-PiNLS-GFP. To obtain pJH2.6-PiNLS-Cas9 and pJH2.6-PiNLS-Cas9-GFP, the NLS region from PITG_11668 was inserted into the SacII and SpeI sites of pYF2.2-PsNLS-Cas9 and pYF2.2-PsNLS-Cas9-GFP. ΔAvr1 was constructed by overlap extension PCR using primers Avr1_del_F and Avr1_del_R. All primers used in this study are listed in **Table S1**.

### Cas9 *in vitro* activity assay

To obtain template DNA for *in vitro* transcription (IVT), a PCR was performed with a T7 promoter-fused forward primer (marked with extension _pT7_F) and the primer sgRNA_Col_R, using plasmid DNA containing the respective gRNAs as template. Next, gRNA was transcribed with T7 RNA polymerase using MEGAshortscript T7 kit (Thermo Fischer Scientific). IVT was allowed to proceed for 4 h, after which RNA was purified by phenol/chloroform and ethanol precipitation, and analyzed on agarose gel. Target DNA was amplified using the respective primers for full-length PCR products of *Avr1, PiTubA2*, and *PiAP5*. SpCas9 nuclease was purchased (New England Biolabs). The assay was performed according to manufacturer’s instructions.

### Molecular analysis of transformants

Genomic DNA (gDNA) was extracted from pooled or individual *P. infestans* transformants according to methods described previously (Fang & Tyler 2016), with modifications. Pooled transformants, 24-48 h after transformation, were pelleted by centrifugation, resuspended in 500 mL of gDNA extraction buffer (200 mM Tris, pH 8.0, 200 mM NaCl, 25 mM EDTA, pH 8.0, 2% SDS, plus 0.1 mg/mL RNase A added prior to use) and sheared by vigorous pipetting. For individual transformants, approximately 250 µl of *P. infestans* mycelium was frozen in liquid nitrogen, freeze-dried, and ground to a powder using a Retsch Mixer Mill MM 400 and metal beads (ø 3 mm) for 30 seconds at 30 Hz. Subsequently, the powder was resuspended in 500 µl of gDNA extraction buffer. DNA was recovered by phenol/chloroform extraction and isopropanol precipitation. RNA was isolated using home-made TRIzol (Verdonk 2014), and cDNA was synthesized using M-MLV Reverse Transcriptase (Promega) according to manufacturer’s instructions. RT-PCR was performed using primers Cas9_RT_F and Cas9_RT_R.

All PCR amplifications were conducted using Q5 high-fidelity DNA polymerase (New England Biolabs). Nested PCR was performed using diluted (1000x) PCR products as a DNA template. PCR amplicons were purified using the NucleoSpin Gel and PCR Clean-up kit (Macherey-Nagel). Sequence analysis was performed at Eurofins.

### T7EI assay

The efficiency of the CRISPR/Cas9 system was tested with T7 endonuclease I (T7EI) (New England Biolabs), using gDNA isolated from pooled transformants as template. First, the target gene was amplified from gDNA using primers amplifying the full-length sequence of the respective target genes (Table S1). Next, purified amplicons were annealed in a thermocycler using the following conditions: 95°C for 5 min, ramp down to 85°C at −2 °C/s, ramp down to 20°C at −0.1 °C/s, hold at room temperature. The T7EI cleavage assay was performed according to manufacturer’s instructions and analyzed by gel electrophoresis.

### RNP assembly and transfection

Ribonucleoprotein complexes (RNPs) of Cas9 (EnGen Cas9 NLS, New England Biolabs) and gRNAs were prepared immediately before transfections. After IVT, gRNAs were refolded by heating at 90°C for 5 min and cooling to room temperature over the course of 12 h. To preassemble RNPs, equimolar amounts of Cas9 (120 pmol) and gRNA (120 pmol) were incubated at 25°C for 10 min. Next, RNPs (10 µl) were incubated with an equal volume of lipofectin for 5 min at room temperature and added to a 100 µl protoplast suspension (2 × 10^6^/ml). After 5 minutes, 120 µl freshly prepared PEG solution (50% PEG-3350, 10 mM CaCl_2_, 10 mM Tris-HCl pH7.5) was added slowly. After incubation for 5 min at room temperature the volume was adjusted to 10 ml with regeneration medium (RSM + 1 M mannitol, without antibiotics) and protoplasts were regenerated for 48 h at 22°C. Next, regenerated protoplasts were collected by centrifugation, from which a sample was taken for gDNA extraction, while the remainder was plated on RSA plates. Protoplasts co-transfected with a HDR construct were plated on RSA plates supplemented with 25 mg/ml hygromycin-B. Colonies appeared on non-selective plates after two days, or after four days on selective plates. Individual colonies were transferred to new plates as soon as they appeared and cultured for further analysis.

## Supplementary files

**Figure S1.**
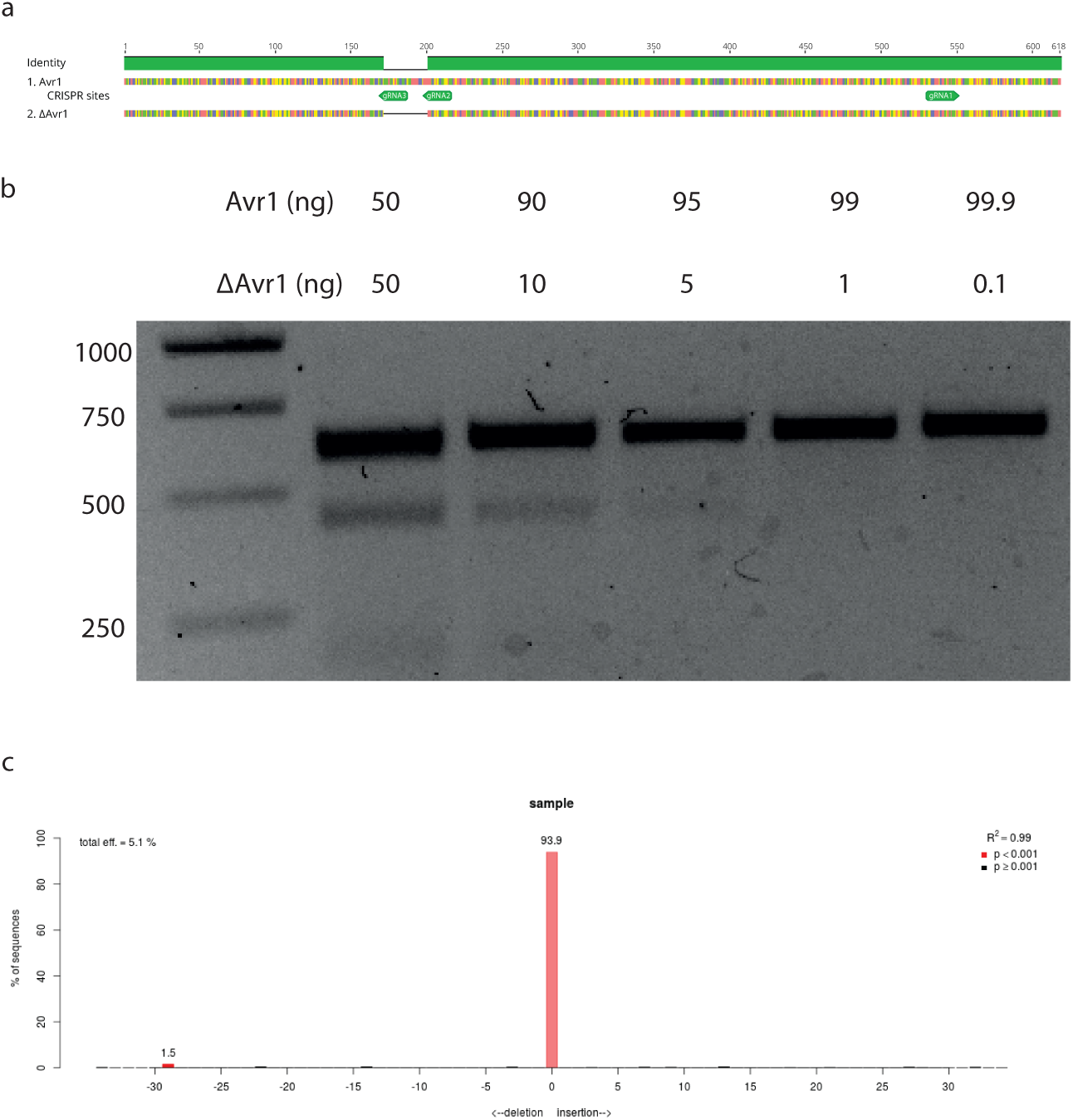
Detection limit assays. **a)** PCR amplicons used for the detection limit assays. ΔAvr1 contains a 29 bp deletion. **b)** T7EI assay. Varying amounts of Avr1 and ΔAvr1 PCR amplicons were annealed and digested by T7EI. **c)** Example output from TIDE, confirming the presence of the 29 bp deletion in ΔAvr1 (left red bar). Here, a sequence chromatogram obtained from sequencing a molar ratio of 998:2 Avr1:ΔAvr1 was compared to a sequence chromatogram of Avr1. The predicted frequency of chromatograms with a deletion (1.5%) deviates from the actual molar ratio of Avr1:ΔAvr1 (0.2%).

**Figure S2.**
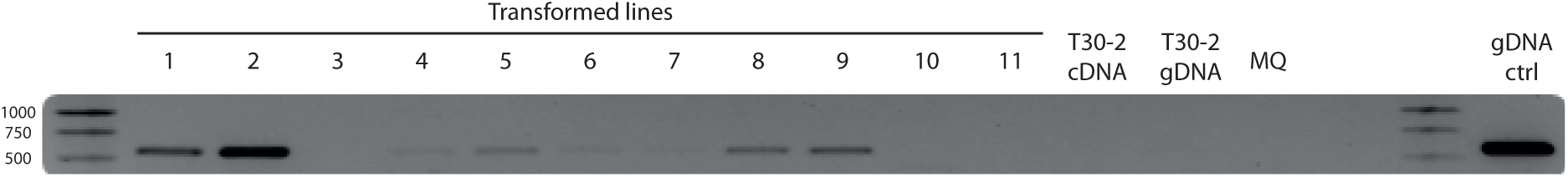
Expression analysis of Cas9 in 11 selected transformed lines. cDNA and genomic DNA (gDNA) of *P. infestans* strain T30-2 were used as negative control and gDNA from a *P. infestans* Cas9 transformant as positive control.

**Figure S3.**
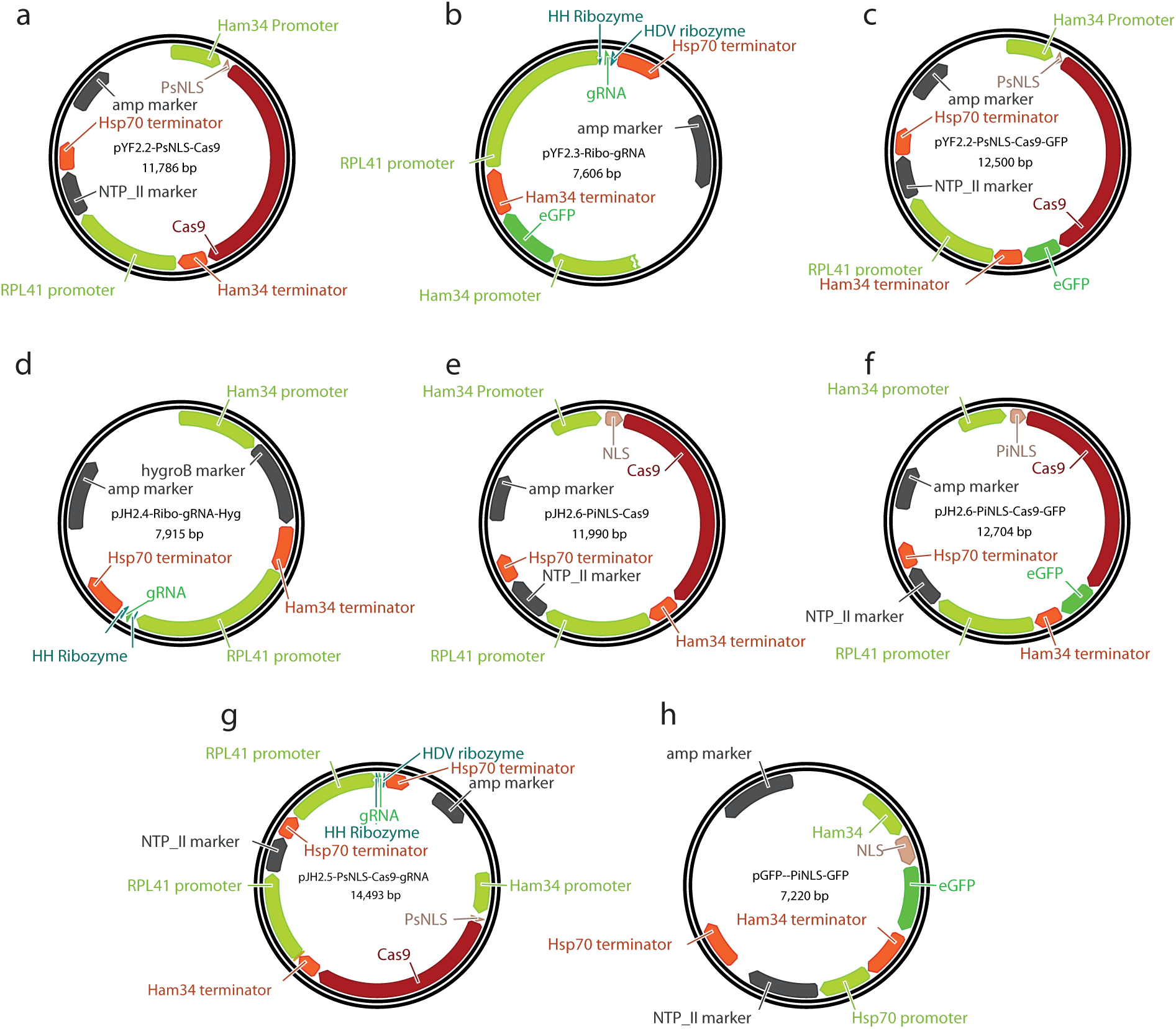
Plasmids used in this study. Plasmids starting with the prefix ‘pYF’ were constructed and kindly provided by Francis Fang (Fang & Tyler 2017), Plasmids with the prefix ‘pJH’ are modified versions as described in this report.

**Table S1.**
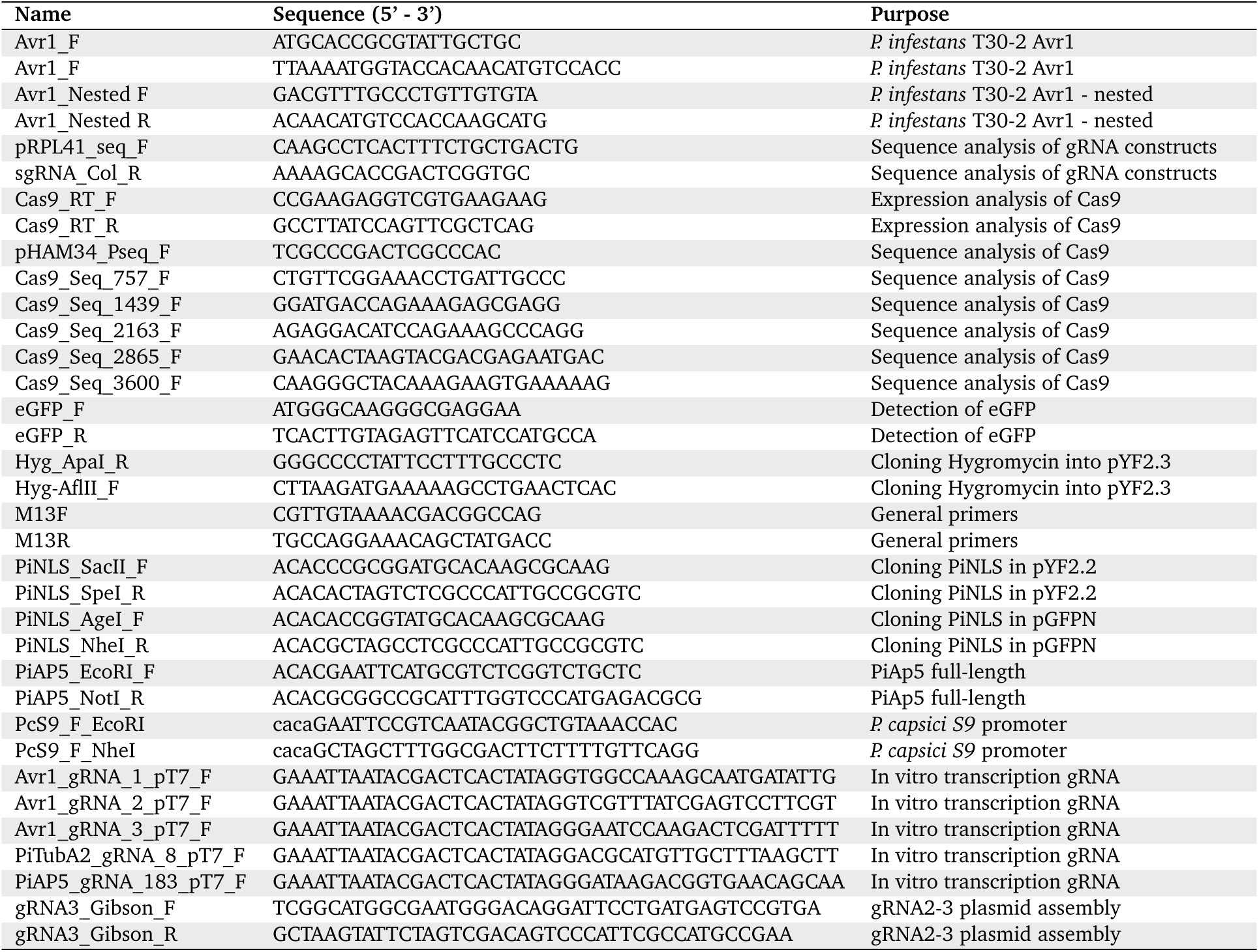
Primers used in this study.

